## Supplementary material for "The antimicrobial peptide Defensin cooperates with Tumour Necrosis Factor to drive tumour cell death in *Drosophila*": Sup Material

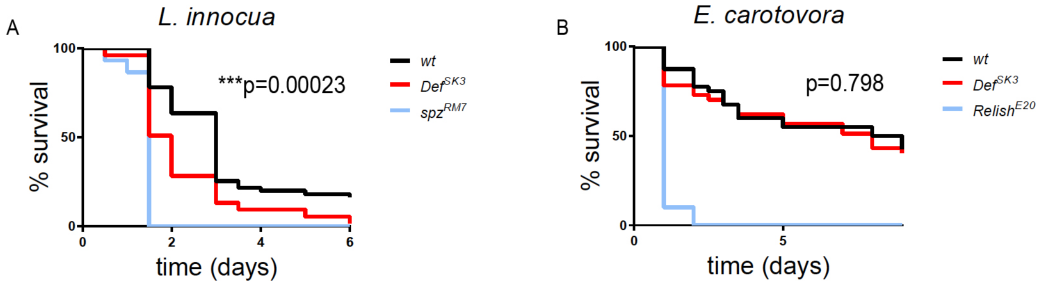


### Figure supplement 1

**Figure supplement 1 | *def* mediates animal survival to infection by Gram-positive bacteria.** A, Survival of *def*$^{sk3}$ mutant flies (n=53), upon Gram-positive, *Listeria innocua* infection, is compared to wild-type (*wt*) (n=55) and *spatzle (spz*$^{M7}$*)* (n=30) (the Toll ligand) mutant flies. B, Survival of *def*$^{sk3}$ mutant flies (n=37), upon Gram-negative, *Erwinia carotovora carotovora* infection, is compared to wild-type (*wt*) (n=40) and *relish (rel*$^{E20}$*)* (n=10) (an Imd-pathway downstream effector) mutant flies. Statistical analysis: A, B, Log-rank test, A, ***p=0.00023, B, p=0.798.

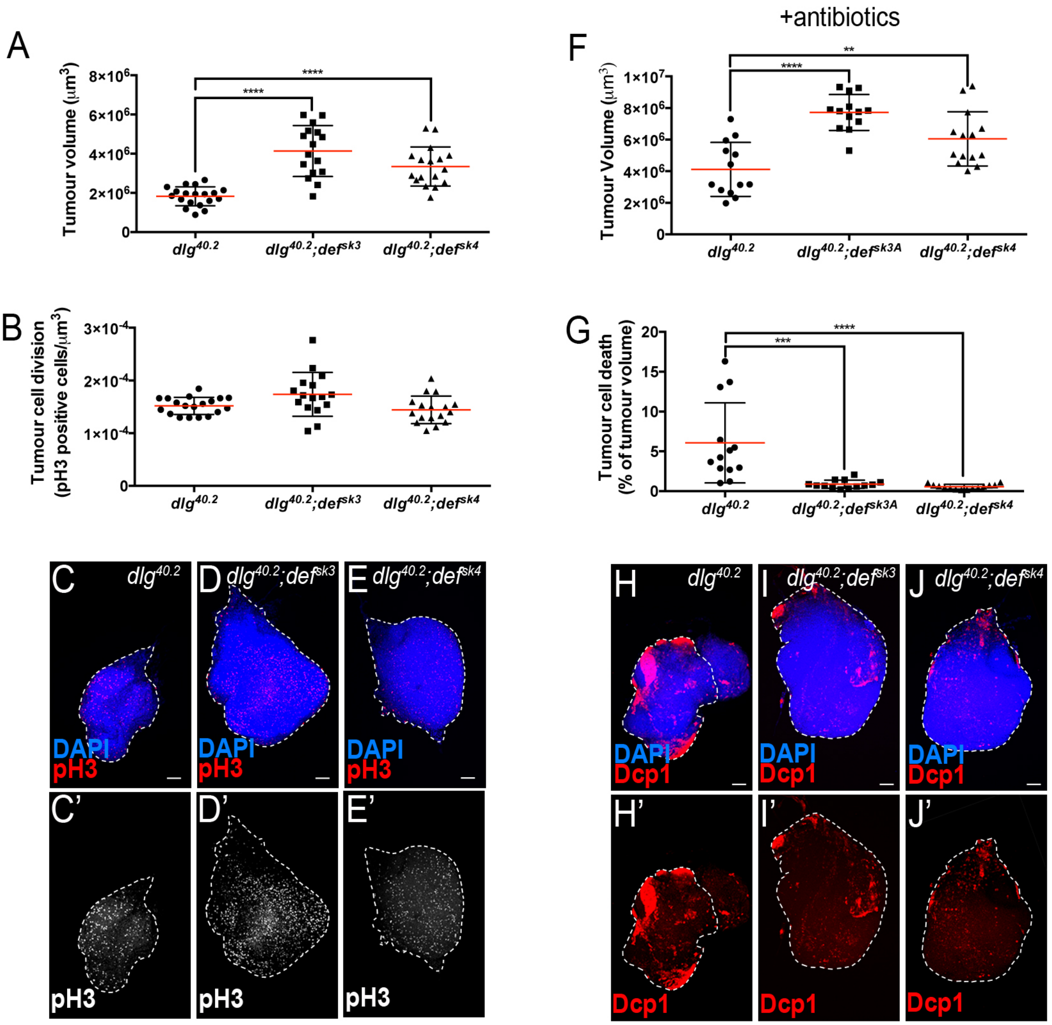


Figure supplement 2

**Figure supplement 2 | Tumour suppression by Def is independent of infection and tumour cell proliferation.** A-E', Quantification of TV (A) and tumour proliferation (B) from *dlg*$^{40.2}$ (n=19), *dlg*$^{40.2}$*;def*$^{sk3}$ (n=16) and *dlg*$^{40.2}$*;def*$^{sk4}$ (n=17) mutant larvae and representative pictures of the corresponding tumours stained with DAPI (blue) or anti-phosphoHistone H3 (pH3) antibody to determine cell proliferation (red, C-E or white, C'-E'). F-J', Quantification of TV (F) and TCD (G) from *dlg*$^{40.2}$ (n=13), *dlg*$^{40.2}$*;def*$^{sk3}$ (n=13) and *dlg*$^{40.2}$*;def*$^{sk4}$ (n=14) mutant larvae reared on antibiotics and representative pictures of the corresponding tumours stained with DAPI (blue) and anti-Dcp1 antibody (red) (H-J'). Scale bars=50 µm. Statistical analysis: A, B, F, G, One way ANOVA, **p<0.01, ***p<0.001, ****p<0.0001.

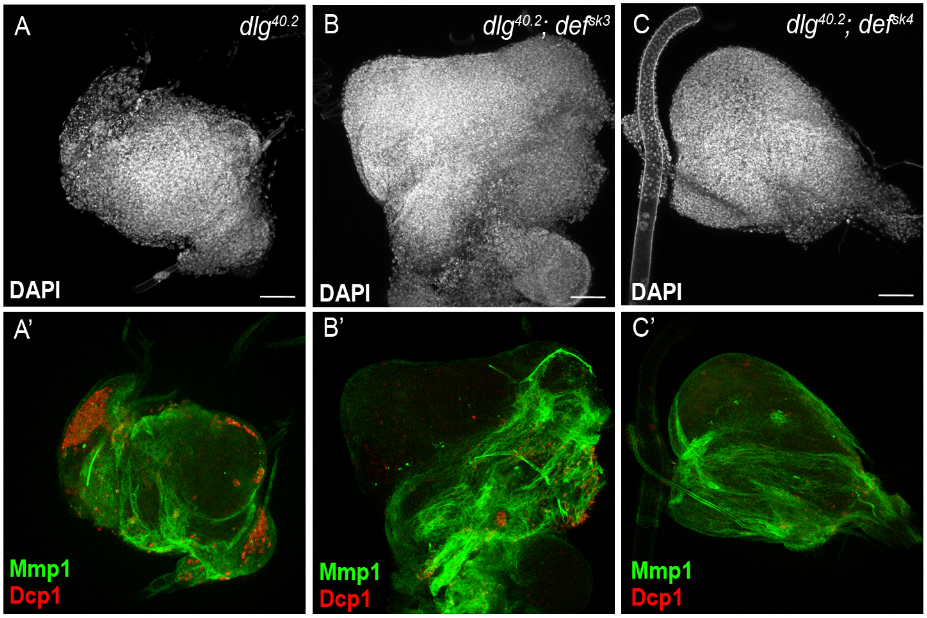


### Figure supplement 3

**Figure supplement 3 | TNF signalling activation does not require a functional *def*.** A-C', DAPI (white), anti-Dcp1 (red) and anti-Matrix metalloproteinase 1 (Mmp1, a reporter of JNK pathway activation, green) antibody staining of wing disc from $dlg^{40.2}$, $dlg^{40.2};def^{sk3}$ and $dlg^{40.2};def^{sk4}$ mutant larvae. Scale bars=50 µm.

**Table supplement 1: List of fly strains used in this work**

| Genotype | Origin | Additional |
|---|---|---|
| $w^{1118}$ | Dewey et al., 2004 | N/A |
| $w^{1118}$ *iso* | Ferreira-Teixeira et al, 2014 | N/A |
| | David Bilder (Mendoza-Topaz et al., 2008) | N/A |
| $egr^{3}$ | Igaki et al., 2002 | N/A |
| $imd^{1}$ | Leulier et al., 2000 | N/A |
| $rel^{E20}$ | Hedengren et al., 1999 | N/A |
| $def^{sk3}$ | This paper | N/A |
| $def^{sk4}$ | This paper | N/A |
| *btl-gal4,UAS-RFP/CyO* | Irene Miguel-Aliaga | N/A |
| *lpp-gal4/TM6B* | Brankatschk and Eaton, 2010 | N/A |
| *tub-gal4/TM3* | BDSC | Cat# 5138 |
| $hml^{\Delta}$*-gal4,UAS-gfp* | Bruno Lemaitre | N/A |
| *en-gal4, UAS-RFP/CyO* | BDSC | Cat# 30564 |
| *UAS-def IR* | VDRC | Cat# FBst0474306 |
| *UAS-imd IR* | VDRC | Cat# FBst0473707 |
| *UAS-myd88 IR* | VDRC | Cat# FBst0455868 |
| *UAS-dlg IR* | VDRC | Cat# FBst0463952 |
| *UAS-egr IR* | VDRC | Cat# FBst0480608 |
| *UAS-w IR* | BDSC | Cat# 25785 |

| | | |
|---|---|---|
| *UAS-def* | Tzou et al, 2002 | N/A |
| *UAS-def-3xHA* | FlyORF | Cat# FBst0501828 |
| *UAS-dcr2* | BDSC | Cat# 24650 |
| *spz*$^{M7}$ | Neyen et al., 2014 | N/A |

**Supplementary Table S2: Primer information**

| **Target gene** | **Primer name** | **Sequence 5’→ 3’** |
|---|---|---|
| *rpl32* | rpl32-fwd | AGGCCCAAGATCGTGAAGAA |
| | rpl32-rev | TGTGCACCAGGAACTTCTTGA |
| *def* (transcript) | def-fwd | CTTCGTTCTCGTGGCTATCG |
| | def-rev | ATCCTCATGCACCAGGACAT |
| *def* (DNA amplification) | def-PCR-fwd | TTATTGCAGAAACGGGCTCT |
| | def-PCR-rev | ATGGTAAGTCGCTAACGCTAATG |
| *def* (DNA sequencing) | def-seq | CGTGTCTTCCTGCACAGAAA |
| *attA* | attA-fwd | ATGCTCGTTTGGATCTGACC |
| | attA-rev | TCAAAGAGGCACCATGACCAG |
| *cecA1* | cecA1-fwd | CTCAGACCTCACTGCAATAT |
| | cecA1-rev | CCAACGCGTTCGATTTTCTT |
| *dro* | dro-fwd | CGTTTTCCTGCTGCTTGCTT |
| | dro-rev | GGCAGCTTGAGTCAGGTGAT |
| *dros* | dros-fwd | CTCTTCGCTGTCCTGATGCT |
| | dros-rev | ACAGGTCTCGTTGTCCCAGA |
